# On the design of treatment schedules that avoid chemotherapeutic resistance

**DOI:** 10.1101/325381

**Authors:** Y. Ma, P.K. Newton

## Abstract

We introduce a method of designing treatment schedules for a model three-component replicator dynamical system that avoids chemotherapeutic resistance by controlling and managing the competitive release of resistant cells in the tumor. We use an evolutionary game theory model with prisoner’s dilemma payoff matrix that governs the competition among healthy cells, chemo-sensitive cells, and chemo-resistant cells and the goal is to control the evolution of chemo-resistance via the competitive release mechanism. The method is based on nonlinear trajectory design and energy transfer methods first introduced in the orbital mechanics literature for Hamiltonian systems. By using the structure of the trajectories defined by solutions of the replicator system for different constant chemotherapeutic concentrations (which produces a curvilinear coordinate system spanning the full region), we construct periodic (closed) orbits by switching the chemo-dose at carefully chosen times and appropriate levels to design schedules that are superior to both maximum tolerated dose (MTD) schedules and low-dose metronomic (LDM) schedules, both of which ultimately lead to fixation of either sensitive cells or resistant cells. By keeping the three sub-populations of cells in competition with each other, neither the sensitive cell population nor the resitant cell population are able to dominate as we balance the populations indefinitely (closed periodic orbits), thereby avoiding fixation of the cancer cell population and re-growth of a resistant tumor. The schedules we design have the feature that they maintain a higher average population fitness than either the MTD or the LDM schedules.

PACS numbers: 87.23.Kg; 87.55.de; 87.19.Xj; 87.19.lr

## I. INTRODUCTION

The development of chemotherapeutic resistance is the primary reason for recurrance of cancer in patients undergoing treatment, and remains one of the primary challenges in the field of oncology [1–3]. As a tumor grows, and even as tumor cells spread throughout the system and metastasis ensues, standard pre-scheduled chemotherapeutic protocols such as maximum tolerated dose (MTD) and low-dose metronomic schedules (LDM) often show early success as the tumor regresses temporarily. A typical chemotherapeutic cycle might involve one strong dose every three weeks, or a dose for 5 consecutive days, followed by a 28 day rest period [4]. After months of fixed periodic cycles, the cancer often recurs and the tumor begins to regrow. Because of the genetic and cellular heterogeneity of a typical tumor [5], instead of killing all of the cancer cells and therby eliminating the tumor, the chemotherapeutic regime actually selects for a resistant phenotype, a phenomenon known as competitive release [6–9]. The diversity of cells within a tumor effectively protects the tumor from single-line or pre-scheduled chemotherapeutic assaults by allowing for elimination of the chemo-sensitive population in order to accomplish the subsequent release of the chemo-resistant population. By reducing the relative fitness of the sensitive cells, chemotherapy acts as the primary mechanism of natural selection [10].

The characterization of a typical tumor as an adaptive landscape made up of competing cells of varying degrees of fitness, which determine growth rates of the various sub-populations is a more accurate characterization of a tumor and suggests an ecological or evolutionary approach [11–19]. If one had access to time-resolved information [20] on the relative balance of the sub-populations of cells making up the tumor, then one could use chemotherapy as a control device (accuator) to keep the sub-populations in balance, competing with each other indefinitely, without any one of the cancerous sub-populations dominating the landscape [6]. Chemotherapy would then be regarded more as a control and maintenance mechanism than a cure [21, 22].

We introduce a method to carry out such a procedure in a three-component replicator system with a time-dependent controller which determines chemotherapeutic dose schedule and concentrations. Although the model system is nonlinear and the control parameter enters as a coefficient in most all of the terms of the three-component cubic nonlinear system (making classic control schemes non-applicable), we use nonlinear trajectory design techniques introduced and developed in an orbital mechanics context [23–27]. In that body of literature, time-dependent controllers are used to design orbit transfers in a Hamiltonian mechanics setting, piecing together partial orbits at different energy levels and switching energies at carefully chosen times, much like classic Homann transfers for satellite control [28]. While the replicator system we describe is not a Hamiltonian system in which orbits transfer from one *energy* level to another, the time-dependent chemotherapeutic parameter, *C*(*t*), can be used to design adventageous orbits in the replicator dynamics tri-linear phase space in a similar manner where piecewise constant dose concentrations are used, *C*_*i*_ = *const*. for carefully chosen time intervals *t*_*i*_ < *t* < *t*_*i*+1_, (*i* = 0,…, *n*) with switching times *t*_*i*_ chosen in such a way as to produce a periodic (closed), continuous, piecewise differentiable orbit that stays trapped in a desirable region of the phase space. Orbits designed this way are shown to maintain a higher average level of fitness for the full population and avoid tumor recurrance. The existence of such orbits for appropriately chosen chemotherapeutic schedules in our model system suggests the possibility that similar orbits may also exist in a more complex tumor environment with carefully designed adaptive schedules [29, 30].

## II. THREE-COMPONENT REPLICATOR SYSTEM

Our model is based on the competitive interactions of three sub-populations of cells, denoted: 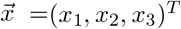. Here, *x*_1_ represents the healthy cells (H), *x*_2_ represents the sensitive cells (S), and *x*_3_ represents the resistant cells (R), with *x*_1_ + *x*_2_ + *x*_3_ = 1. The replicator dynamical system is then:

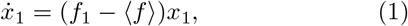

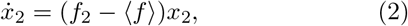

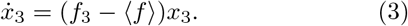

The fitness functionals are given by:

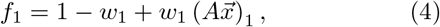

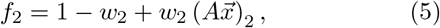

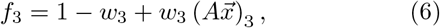

where 0 ≤ *w*_*i*_(*t*) ≤ 1, (*i* = 1, 2, 3) are time-dependent selection parameters (that serve as our controllers) we use to shape the fitness landscape of the system. The time dependence enters through the chemo-concentration parameter *C*(*t*):

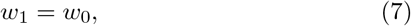

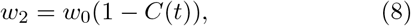

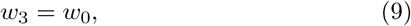

where *w*_0_ scales time (we typically take *w*_0_ = 1). Note that the chemotherapy parameter acts directly only on the sensitive cell population lowering its fitness, although the three populations are coupled nonlinearly through (1)-(3). The average fitness is given by the functional:

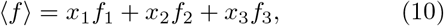

and the 3 × 3 payoff matrix is:

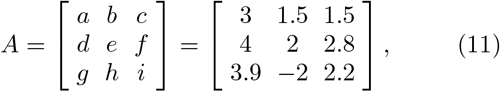

which we take as a prisoner’s dilemma matrix for every two-by-two sub-block [31]:

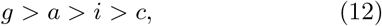

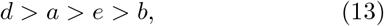

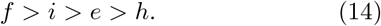

Matrix entries are consistent with those from [32]. As discussed in [32], the prisoner’s dilemma payoff matrix ensures: (i) Gompertzian growth of the cancer cells; (ii) a reduction in overall fitness of the population as the tumor grows; and (iii) a fitness cost associated with resistance. In our previous paper [32] we studied the nonlinear dynamics associated with eqns (1)-(3) for constant values of the chemotherapy parameter 0 ≤ *C* ≤ 1 to demonstrate the mechanism of competitive release when C ≥ 1/3. In this paper, we investigate piecewise constant time-dependent functions *C*(*t*) to show how to avoid the evolution of resistance of the tumor. Figure 1 shows several examples of the chemotherapeutic schedules we consider. These include maximum tolerated dose (MTD) schedules (Figure 1(a)), low-dose metronomic schedules (LDM) (Figure 1(b)), adaptive schedules (Figure 1(c)), and more general time-dependent schedules (Figure 1(d)) which we break-up into piecewise constant doses as done in forming the Riemann sum approximation to an area under a curve. In all cases, we compare outcomes of the different schedules holding the total dosage, *D*, over a fixed time period *τ*,

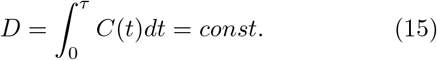

**FIG. 1.**
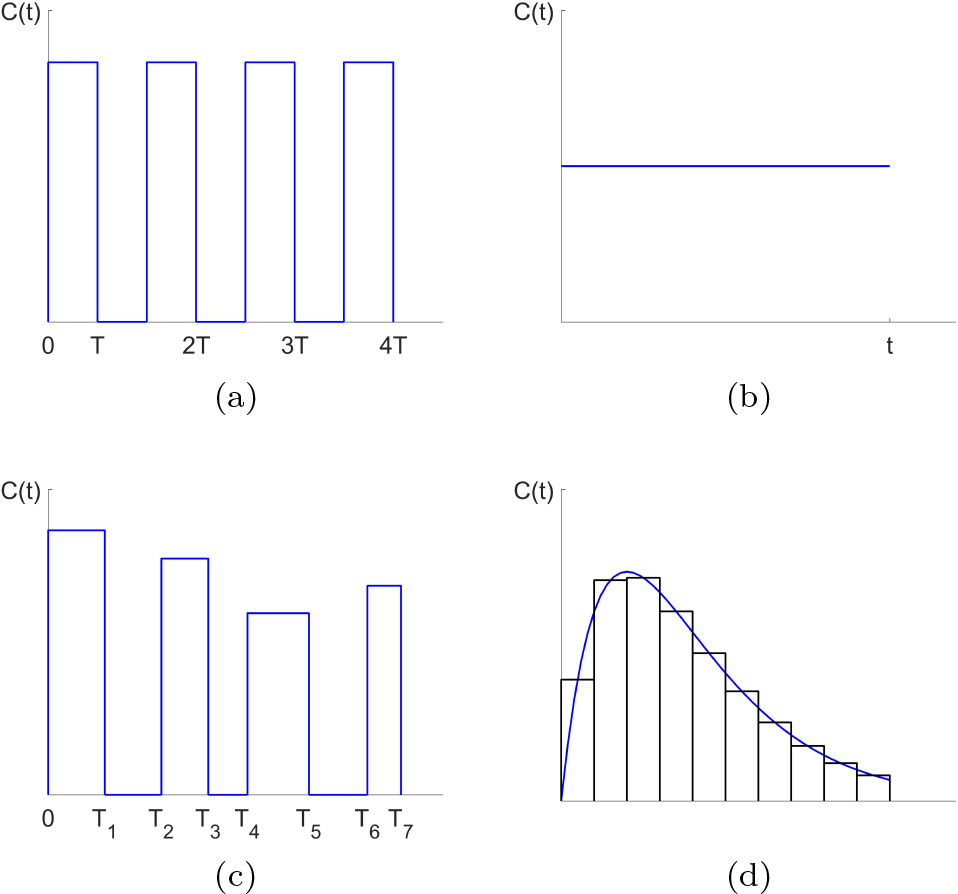
Dose schedules. (a) Maximum tolerated dose (MTD) schedule; (b) Low dose metronomic (LDM) schedule; (c) Adaptive schedule; (d) General time-dependent schedule accuated by piecewise constant dose concentrations.

## III. RESULTS

Figure 2 shows the key idea behind the method we use to design trajectories for eqns (1)-(3). With no chemotherapy, *C* = 0, since the sensitive corner *S* is a globally asymptotically attracting fixed point whose basin of attraction is the full region, all trajectories that start inside the triangle eventually get trapped in the left *S* corner. By contrast, with chemotherapeutic levels at *C* = 0.7, competitive release acts to create a basin of attraction for the right resistant corner *R* for all initial conditions inside the triangle. Using these families of solution trajectories, we overlay the solution curves in Figure 2(c) to show the underlying curvilinear grid that spans the full tri-linear phase plane. By switching between the two values *C* = 0 and *C* = 0.7 at times when two curves intersect, it is possible to transition from a trajectory associated with the *C* = 0 family to one associated with the *C* = 0.7 family. This creates multiple possibilities for designing complex orbits using piecewise constant values of *C*, with switching at appropriately chosen times. One such trajectory is shown in Figure 3(a) achieved by switching between the two values *C* = 0 and *C* = 0.5. The trajectory starts at point *A* ((*H*, *S*, *R*) = (0.5, 0.4, 0.1)) using a trajectory with *C* = 0.5. When the trajectory reaches point *B*, the chemotherapy is switched off *C* = 0, and the system then travels back down to point *A* in a closed loop. With switching times labeled *T*_1_ and *T*_2_, this closed orbit can be maintained indefinitely. By contrast, Figure 3(b) shows the corresponding MTD and LDM trajectories with the same initial condition (point *A*) using the same total dosages over the same time-periods. The MTD trajectory eventually gets trapped in the left *S* corner, as does the LDM trajectory. Figure 3(c) shows the schedules for all three cases. In all cases, we first design the adaptive schedule, the we create the MTD and LDM schedules using the same total dosage *D*. Neither the MTD nor the LDM standard chemo-schedules is able to prevent the system from saturating with a full grown tumor, whereas the adaptive schedule keeps the system trapped indefinitely near the top *H* corner of the system

**FIG. 2.**
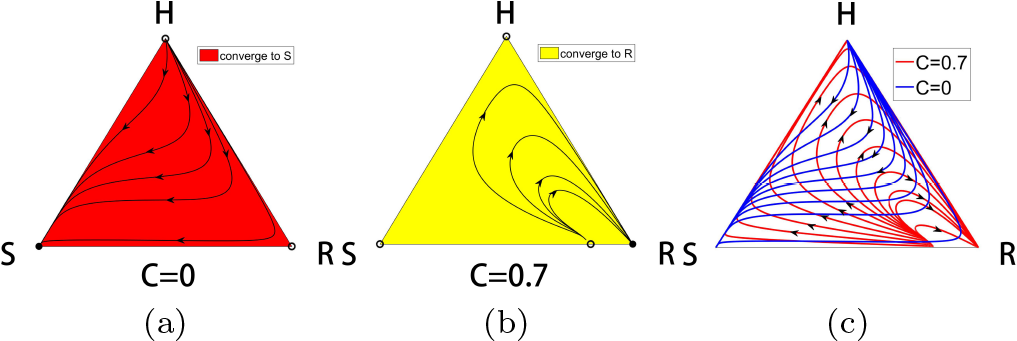
Dynamical trajectories for constant *C*. (a) With no chemotherapy (*C* = 0), the sensitive corner *S* is a globally attracting fixed point. All initial conditions inside the triangle move to *S* along the sample trajectories shown. (b) Above the chemotherapy threshold *C* > 0.5, all initial conditions inside the triangle move to the resistant corner *R*. Shown are sample trajectories for *C* = 0.7. (c) Overlay of the solution trajectories for *C* = 0 and *C* = 0.7 create a curvilinear grid throughout the triangle. By switching between these two values of *C*, we construct a global trajectory made up of segments of the two families of trajectories.

**FIG. 3.**
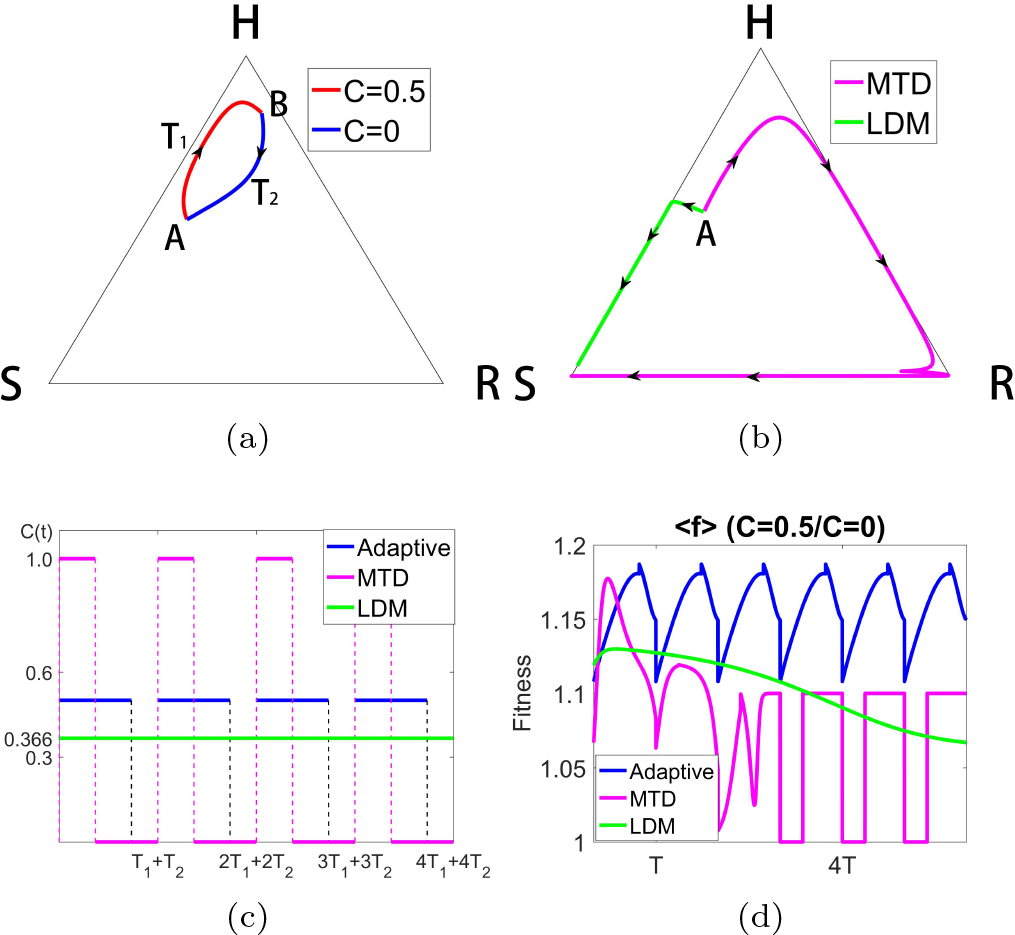
Constructing a closed loop trajectory using *C* = 0 and *C* = 0.5, with *C*_*avg*_ = 0.366. (a) By using segments of a trajectory for *C* = 0 and *C* = 0.5, switching values at points *A* and *B*, we construct a closed periodic orbit. (b) Using the same total dose, we show the MTD and LDM trajectories starting from point *A*. Both eventually move to the *S* corner, although initially the MTD trajectory moves toward the *H* corner (tumor regression) before recurrance. (c) The MTD, LDM, and adaptive schedules are depicted. (d) We plot the average fitness 〈*f*〉 for the three different chemo-schedules. The adaptive schedule is able to maintain a higher average fitness throughout.

Figure 3(d) shows the average fitness of the system for the MTD, LDM, and adaptive schedules. It is clear that the adaptive schedule is able to maintain a higher average fitness throughout the full course of chemotherapy.

Figure 4 shows the sensitivity of the system to the chemo-concentration levels chosen. Here we switch between *C* = 0 and *C* = 0.6 (higher average dose than in Figure 3) to construct the closed loop (Figure 4(a)), which looks very similar to that in Figure 3. But notice that for these values, the LDM schedule creates an orbit that saturates at the *R* corner, whereas the MTD schedule saturates at the *S* corner. The actual schedules are shown in Figure 4(c). Figure 4(d) compares the average population fitness of the three schedules, showing the adaptive schedule maintaines a higher average throughout. Notice that LDM initially achieves a higher average fitness before tumor regression occurs.

**FIG. 4.**
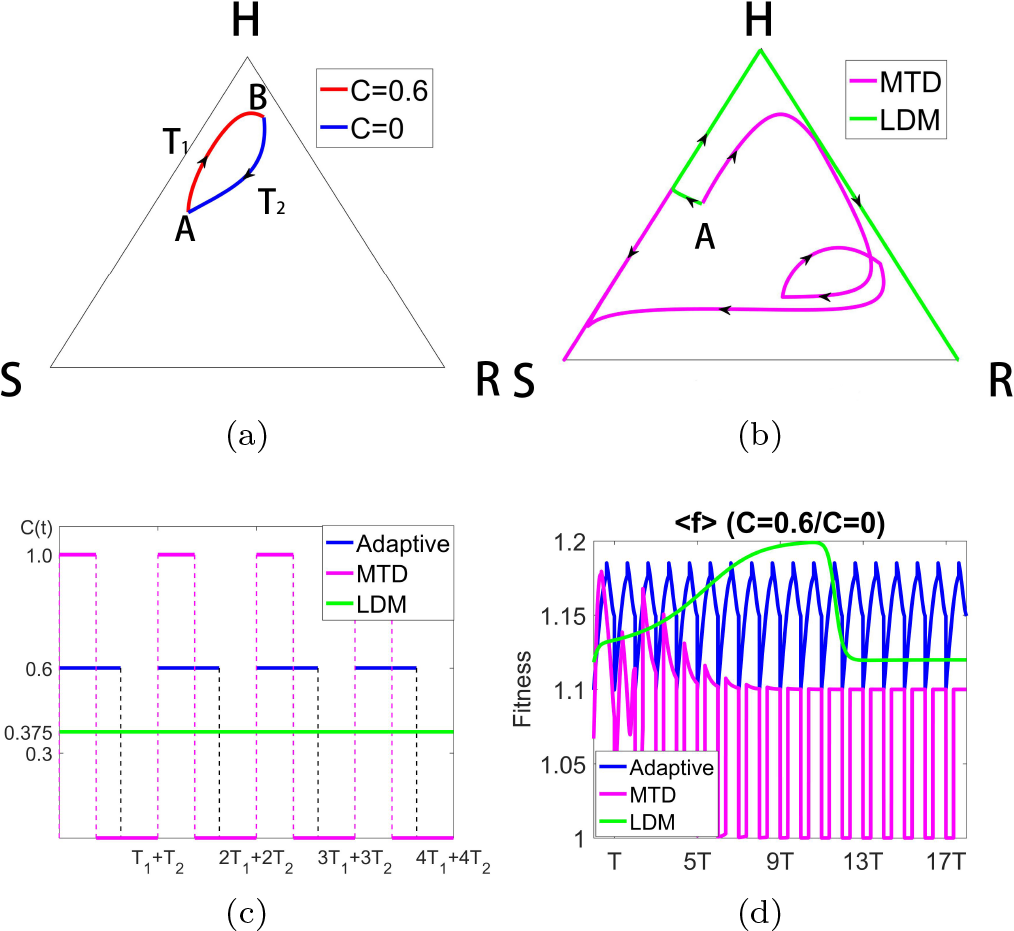
Constructing a closed loop trajectory using *C* = 0 and *C* = 0.6, with *C*_*avg*_ = 0.375. (a) By using segments of a trajectory for *C* = 0 and *C* = 0.6, switching values at points *A* and *B*, we construct a closed periodic orbit. (b) Using the same total dose, we show the MTD and LDM trajectories starting from point *A*. The MTD trajectory eventually moves to the *S* corner, while the LDM trajectory moves to the *R* corner. Both trajectories initially move toward the *R* corner (tumor regression) before recurrance. (c) The MTD, LDM, and adaptive schedules are depicted. (d) We plot the average fitness 〈 *f* 〉 for the three different chemo-schedules. The adaptive schedule is able to maintain a higher average fitness throughout, although LDM initially achieves higher average fitness before declining.

In Figure 5 we show the result of toggeling between values *C* = 0.3 and *C* = 0.6 to maintain the periodic loop (Figure 5(a)). For this case, both the MTD and LDM schedules send the trajectory to the *R* corner (Figure 5(b)). Figure 5(d) shows the initial benefit of the MTD and LDM schedules in terms of higher initial average fitness, but eventually the adaptive schedule shows its superiority over both.

**FIG. 5.**
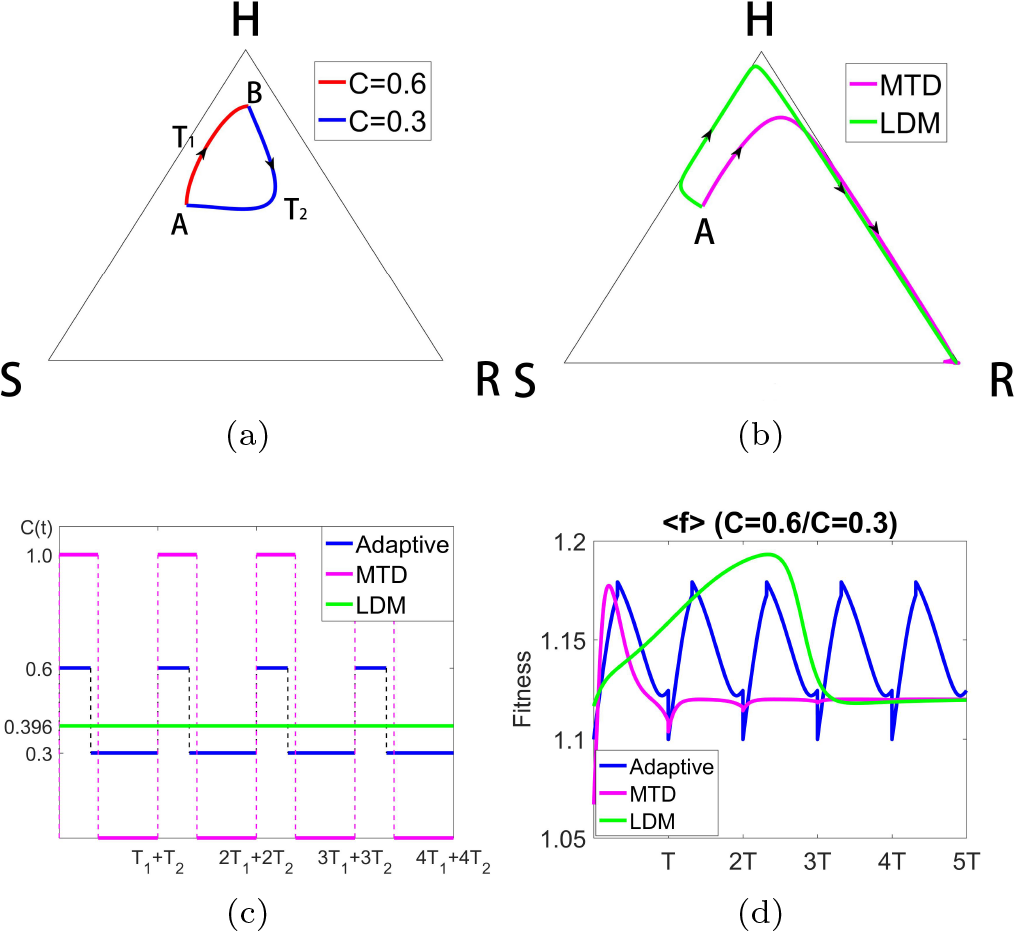
Constructing a closed loop trajectory using *C* = 0.3 and *C* = 0.6, with *C*_*avg*_ = 0.396. (a) By using segments of a trajectory for *C* = 0.3 and *C* = 0.6, switching values at points *A* and *B*, we construct a closed periodic orbit. (b) Using the same total dose, we show the MTD and LDM trajectories starting from point *A*. Both trajectories move toward the *R* corner after initially moving toward *H*. (c) The MTD, LDM, and adaptive schedules are depicted. (d) We plot the average fitness 〈*f*〉 for the three different chemo-schedules. The adaptive schedule is able to maintain a higher average fitness throughout. Both MTD and LDM initially show higher average fitness before declining.

Figure 6(a) shows that we can actually design an orbit that starts at an arbitrary point *A* inside the triangle, and send it to an arbitrary point *B*. We accomplish this by constructing the incoming and outgoing orbits from point *A* for two different *C* values, and those associated with point *B* for those same two *C* values, showing that they must intersect at some point which we label *O*. By sending the orbit out from point *A* to point *O*, then switching values of *C* at point *O* until we arrive at point *B*, we complete the transfer. One can immediately see the potential richness in the possible design of different orbits one can construct by switching values of *C* among two, three, or more values, at appropriately chosen times. Figure 6(b) shows the richness of the curvilinear grid that can be created with three values of *C* = 0,0.3,0.7 and the multitude of possible paths from one point to another inside the triangle if one allows for switching among the three values shown.

**FIG. 6.**
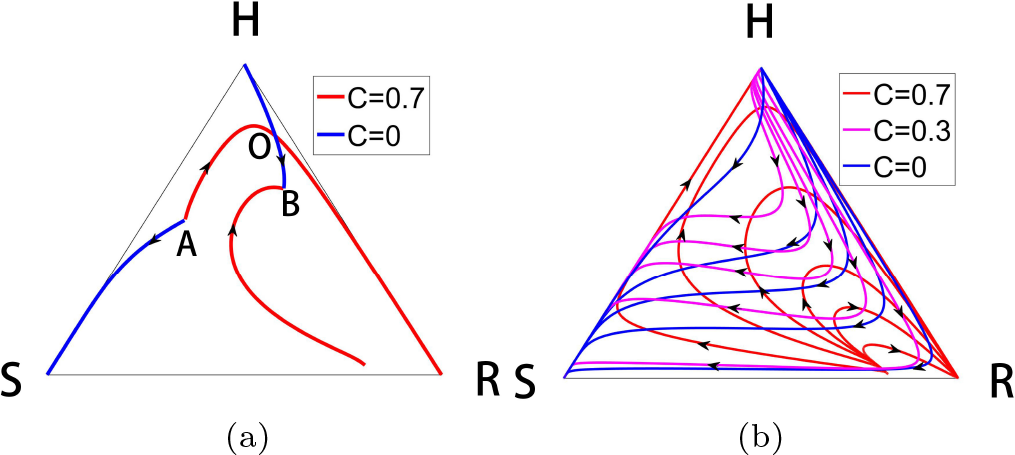
Constructing orbit transfers. (a) We take two arbitrary points labeled *A* and *B* and construct the incoming and outgoing solution trajectories from each, using two values *C* = 0 and *C* = 0.7. There must be a crossing point, which we label point *O*. The two segments *AO* followed by *OB* is the two-switch trajectory that takes us from *A* to *B*. (b) Shown is the curvilinear grid constructed by families of solution curves for the three values *C* = 0, *C* = 0.3, *C* = 0.7. Any point of intersection can be used as a starting point and an ending point to construct a three-switch orbit.

## IV. DISCUSSION

One of the challenges of designing optimized schedules with clinical trials is that short term gain in average fitness with LDM or MTD does not always result in more long-term increased averaged fitness levels, i.e. recurrance sets in if the schedule does not completely eradicate the tumor. This is clearly shown in our simulations. The possibility of designing complex orbits with various potentially adventageous features is virtually endless if one allows for enough switches among many different levels of the chemo-concentration parameter *C*. In this manuscript, there has been no attempt at building optimality into any of the designed orbits, aside from a comparison of average fitness levels. One could, of course, imagine designing orbits that maximize a (time-averaged) population fitness while minimizing total dose, or perhaps delay recurrance as long as possible while avoiding regions of the phase space (i.e. relative balances of the sub-population levels) where the total tumor burden becomes unsustainable. The fact that closed loop trajectories can be designed in our model three-component replicator system simply via chemo-scheduling indicates the possibility that similar orbits could potentially be created in an actual tumor environment, with the right accuation (chemo-schedule), where fixation of the sensitive population or fixation of the resistant population are both avoided indefinitely as the balance is managed with proper adaptive therapy. This, of course, all hinges on our ability to carefully and frequently monitor the tumor environment [20]. Clinical trials that make use of adaptive scheduling ideas are currently being carried out at the Moffitt Cancer Center, and results show great promise [29]. More possibilities exist with the use of combination multi-drug therapies [13], or immunotherapies [33], but these are outside the scope of our model.

## Acknowledgments

We gratefully acknowledge partial support from the Breast Cancer Research Foundation (BCRF) and the Jayne Koskinas & Ted Giovanis Foundation (JKTG) for Health and Policy. We thank S. Anderson, D. Basanta, J. Brown, H. Enderling, R. Gatenby, and J. West at the Moffitt Cancer Research Center for helpful conversations regarding this and related work.

